# Dense reconstruction of elongated cell lineages: overcoming suboptimum lineage encoding and sparse cell sampling

**DOI:** 10.1101/2020.07.27.223321

**Authors:** Ken Sugino, Rosa L. Miyares, Isabel Espinosa-Medina, Hui-Min Chen, Christopher J Potter, Tzumin Lee

## Abstract

Acquiring both lineage and cell-type information during brain development could elucidate transcriptional programs underling neuronal diversification. This is now feasible with single-cell RNA-seq combined with CRISPR-based lineage tracing, which generates genetic barcodes with cumulative CRISPR edits. This technique has not yet been optimized to deliver high-resolution lineage reconstruction of protracted lineages. *Drosophila* neuronal lineages are an ideal model to consider, as multiple lineages have been morphologically mapped at single-cell resolution. Here we find the parameter ranges required to encode a representative neuronal lineage emanating from 100 stem cell divisions. We derive the optimum editing rate to be inversely proportional to lineage depth, enabling encoding to persist across lineage progression. Further, we experimentally determine the editing rates of a Cas9-deaminase in cycling neural stem cells, finding near ideal rates to map elongated *Drosophila* neuronal lineages. Moreover, we propose and evaluate strategies to separate recurring cell-types for lineage reconstruction. Finally, we present a simple method to combine multiple experiments, which permits dense reconstruction of protracted cell lineages despite suboptimum lineage encoding and sparse cell sampling.

## Introduction

Understanding the complexity of the brain is one of the last great frontiers of biological science. Brain complexity is realized, in part, by a rich diversity of neuronal cell types (for example [1]). The diversity of neuronal types is laid down during the developmental process. One way to help identify mechanisms that establish this diversity is to connect neuronal gene expression with the neuron’s developmental history. Therefore, methods combining transcriptome profiling and lineage tracing have the potential to reveal how neuronal stem cell identity and neuronal birth order are translated into cell-type information.

CRISPR-based lineage tracing combined with single cell RNA-seq is a promising candidate for this purpose. Lineage tracing techniques have evolved from visually tracking lineage development[2, 3, 4, 5], to genetically encoded makers or barcodes used to identify cells in a lineage[6, 7, 8, 9], to CRISPR-based dynamic lineage tracing[10, 11, 12, 13, 14, 15, 16, 17]. Compared to previous methods, CRISPR-based dynamic lineage tracing has the possibility of obtaining a more detailed picture of lineage development. The CRISPR-based method creates evolving genetic barcodes via a cumulative process of CRISPR editing. Cas9, in combination with a gRNA, modifies specific predefined sequences in the cell (for more details see [18, 19, 20]). This editing process (encoding) produces independent edits in different cells. These modifications (barcodes) accumulate during development, thus providing information on cellular lineage. At some point, the cells can be harvested and the barcodes (either DNA or expressed as mRNA) can be sequenced. The lineage relationship of the cells can then be reconstructed using phylogenetic analysis, since similarity of barcodes indicates relatedness.

Our goal is to adapt this CRISPR-based technique for a high-resolution reconstruction of brain development. To accomplish this objective, we chose to start with *Drosophila* type 1 central brain neuroblast lineages as a model system. *Drosophila* type I neuroblasts are extremely well-characterized neural stem cells[21, 22]. Cycling through asymmetric divisions, a typical type I neuroblast yields a deep linear series of ~100 intermediate precursors (ganglion mother cells; GMC) that each divide once, producing a total of about 200 neurons. Early embryonic patterning confers each neuroblast with a unique identity that governs production of lineage-specific neuron types[23]. Exquisite temporal patterning further allows a neuroblast to yield multiple neuron types in an invariant sequence[24, 25]. The birth order and adult neuron morphology has been analyzed at single cell resolution for a number of lineages. Morphological lineage tracing has revealed that lineages can vary greatly in the number of morphologically distinct cell types. For example, in the mushroom body (MB) lineage, there are only three main cell types born consecutively over long stretches[26], whereas in the antennal lobe (AL) lineages, the temporal fate seems to change every two cell cycles, generating ~50 serial neuron types[27]. Recently, we also observed the recurrence of morphological features in separate temporal windows[28]. Together, these features make the well-characterized type 1 neuroblast lineages ideal models to explore cell-type generation.

Despite the potential for detailed, complete lineage reconstruction, early CRISPR-based dynamic lineage tracing methods have not been applied for elongated lineages such as *Drosophila* type 1 neuroblast lineages. The first example of CRISPR dynamic lineage tracing, GESTALT[10], used an array of 10 coding units and provided Cas9 and gRNAs at the beginning of embryonic development. The 10-unit array yielded enormous barcode diversity but suffered from limited duration that it can track due to transient Cas9 activity and rapid coding unit exhaustion. Labs have since worked to increase the encoding capacity by a number of techniques[16, 13, 17, 29]. Studies have also attempted to extend the duration of the encoding process. One lab did this by expressing Cas9 and specific gRNAs both at the beginning of embryogenesis and in later stages of brain development[15]. This study uncovered lineage origins of multi-potent neuronal progenitors, however the depth of the lineage tracing (i.e. how many distinct sequential cell divisions it can track) was insufficient to provide information on how these progenitors yield different neuronal types. Another study designed gRNAs with a range of Cas9 cutting rates; some were fully complementary to the target sites and some had 1-2 mismatches to reduce the editing rate[17], this significantly increased the depth of lineage tracing. However, none of the previous studies targeted lineage as deep as *Drosophila* type 1 neuroblast lineages, therefore it is unclear whether these methods/settings would be effective in tracing deep elongated type 1 lineages.

In this article, we use simulations to first elucidate the theoretical requirements on experimental parameters required to trace deep lineages such as *Drosophila* type 1 neuroblast lineages. We find there are two critical parameters which need to be adjusted depending on the depth of the target lineage. The first is the rate of encoding (Cas9 editing rate). We determine the optimum rate (*r*) to be inversely proportional to the depth of the lineage to be encoded *r* = 1/*depth*. In dividing neuroblasts, we experimentally confirm that the editing rate of an available Cas9 base-editor is very close to the optimum rate for type 1 lineages. The second critical parameter is the number of coding units. The number of targets required to robustly track a lineage with a depth of 100 is more than 1000. As this requirement is experimentally impractical, we examine ways to overcome target number deficiencies. Consequently, we devise a method to combine information from multiple experiments. Merging multiple experiments significantly reduces the required number of coding units. In addition, this also addresses the problem of cell and/or barcode loss (e.g. 15.8-73.7% in Chan et al[17]), which severely hinders the performance of lineage reconstruction.

To combine multiple experiments, we utilize the linear topology of *Drosophila* type 1 lineages. We convert the number of edits in each cell to a pair-wise ordering of cells (order matrix). Due to inadequate numbers of coding units and to cell/barcode loss, each experiment’s order matrix is incomplete or erroneous. However, assuming that cell types can be matched across experiments, these incomplete matrices can be combined to yield a more complete order matrix. From this combined order matrix, we can recover a comprehensive lineage tree.

In summary, in this report, we clarify experimental parameter requirements to trace elongated *Drosophila* type 1 neuroblast lineages. In addition, we propose a method to combine information from multiple experiments, a critical element which has been lacking in this field.

## Results

### The optimum editing rate to encode a typical Drosophila neuronal lineage is 1% per cell cycle

In order to clarify the requirements for CRISPR-encoded lineage tracing of a typical *Drosophila* type 1 neuroblast lineage (100 asymmetric stem cell divisions), we need to first model the dynamics of CRISPR editing. Cas9 nuclease is currently the most commonly used enzyme for CRISPR-encoded lineage tracing, as it creates incredibly diverse edit outcomes or alleles[10]. However inter-target deletions can occur if Cas9 targets are on the same chromosome. It has been shown that frequent, large, inter-target deletions diminish the ability to accurately perform phylogenetic analyses[30, 18]. Inter-target deletions would be particularly problematic if the Cas9 targets are placed in an array, as coding units or previous edits can easily be lost[10]. Inter-target deletions should also be avoided if the number of chromosomes for integration is limited, such as in *Drosophila*. We therefore decided to use a Cas9 base-editor in our modeling. The targets for Cas9 base-editors can be designed to have multiple cytosines, achieving various possible outcomes, or alleles/per unit (see Discussion).

We model the process of editing *nU* units/targets where the rate of editing per unit, per cell division is *r*. For the purposes of modeling, we assume the editing rate is equal and independent across units and over the course of development (Fig.1A). Importantly, once a unit is edited, it cannot be edited again. We denote the number of editing outcomes as *nL* (number of levels/alleles). Here, we set *nL* as two, as this is readily achievable with a Cas9 base-editor. We also give equal probability (*p* = 1/*nL*) of either level being chosen at each editing event. Most of these assumptions are simplified. For example, allele choice is likely to be biased and the editing rate most likely varies between targets and across development. However, we begin with this simplified model to delineate the basic dynamics of CRISPR-barcoding with relatively simple mathematics.

**Figure 1:**
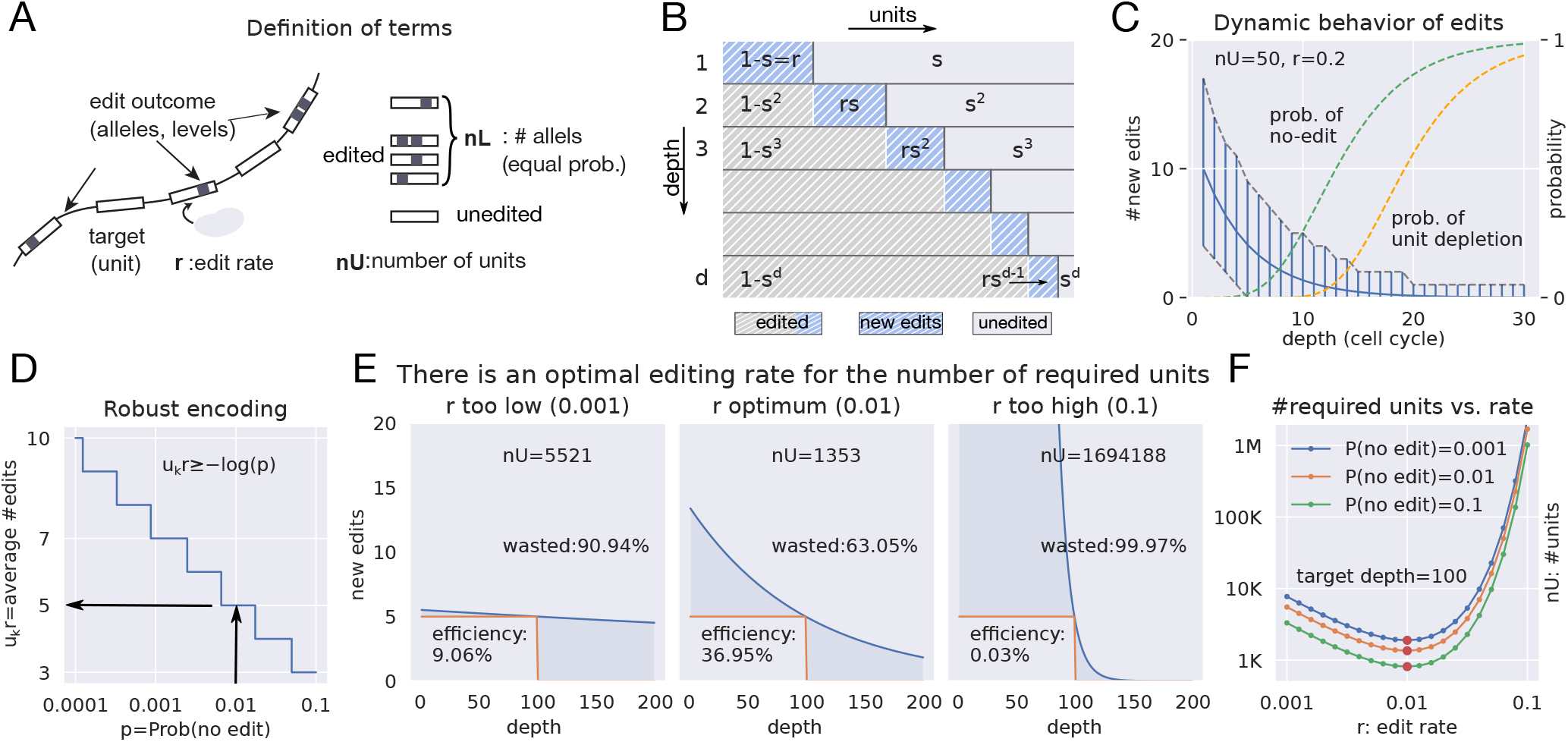
Dynamics of encoding the lineage barcode **A:** Schematics of the model of the CRISPR encoding process. An array of Cas9 targets is shown on the left. Edits are denoted as black mark(s) within a target. Critical parameters: *nU*: number of encoding units, *nL*: number of edit outcomes (alleles), *r*: editing rate (per unit, per cell cycle). Four editing outcome (alleles) are diagramed, although the modeling conservatively uses *nL* = 2. **B:** Average numbers of edited units transition as the lineage progresses (as depth increases). *s* = 1 − *r* is the probability of a unit not being edited (solid gray) during a given cell cycle. As the number of units available for editing decreases, fewer new edits (blue dashes) occur. *d* refers to the final depth of the lineage. **C:** The average number of new edits in successive cell cycles (*k*) decreases exponentially as the probability of no edits and probability of unit depletion increase. Average new edits (*n_k_* = *nUrs*^*k*−1^, solid blue line) and 99% range of new edits (calculated from binomial distribution *B*(*nU*, *rs*^*k*−1^) at each cell cycle, vertical blue lines), probability of no edits (dashed green line, (1 − *rs*^*k*−1^)^*nU*^) and unit depletion (dashed orange line, (1 − *s^k^*)^*nU*^) are plotted against the depth. *nU* = 50 and *r* = 0.2. **D:** Number of average edits required for robust encoding. The probability of no edits (which is 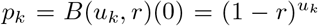, where *u_k_* are number of remaining units at cycle *k*) has to be smaller than the specified confidence level. When *r* is small, −*log*(*p_k_*) = − *u_k_log*(1 − *r*) ~ *u_k_r* is the expected number of edits. Arrows indicate that the average number of edits needs to be five to have less than 1% probability of no edits. **E:** Schematics showing the efficiency *e* for three editing rates for the exponential encoding process. The Y-axis is the number of new edits at each cycle. The shadowed areas indicate wasted units, including redundantly edited units (more than 5 in each cell cycle) and units that remain unused beyond the desired depth (100). If the editing rate is too low (left panel), the waste primarily resides after the 100th cell cycle. If the editing rate is too high (right panel), the waste mainly results from redundant edits during the initial phase. **F:** Effects of editing rates (X-axis) on the number of required units (Y-axis) for various confidence levels (per probability of no edit at the 100th cell cycle). The optimum rate (the rate which uses the units most efficiently) to encode a lineage of depth *d* is 1/*d*, regardless of the probability of no edits. The optimum rate of 1/*d* is derived from taking a derivative of new edits at depth *d*: *n_d_* = *nUr*(1 − *r*)^*d*−1^, requiring the inital number of units *nU* to be minimum (*d*(*n_d_/nU*)/*dr* = (1 − *r*)^*d*−1^ + *r*(*d* − 1)(1 − *r*)^*d*−2^ = (1 − *r*)^*d*−2^(1 − *rd*)). The asymmetry of curved lines demonstrates the need for copious units if edited at higher-than-optimum rates.

For high-resolution reconstruction of a lineage, it is critical to have new edits at each cell cycle. We refer to the total number of serial cell cycles that the founder cell will undergo in a protracted linear lineage as its depth and use *k* to represent an intermediate cell cycle along the depth. As the lineage progresses and units are edited, the number of available units decreases. This therefore decreases the probability of new edits as the lineage progresses (as *k* increases). At a given cell cycle, under the above assumptions, the number of newly edited units follows a binomial distribution with the success probability equal to the rate (*r*) and the number of trials being the number of remaining unedited units (*u_k_*) (Fig.1B). On average, the number of remaining unedited units decreases exponentially: *u_k_* = *nU* (1 − *r*)^*k*^ for a depth of *k*; thus the average expected number of newly edited units (*n_k_*) also drops exponentially: *n_k_* = *nUr*(1 − *r*)^*k*−1^ (Fig.1C). This indicates that the ability to tell ancestral relationships in a lineage also rapidly decreases.

To distinguish consecutively born cells, it is necessary to have some new edits at each cell cycle. To keep the probability of having no edits below 1% (in other words to keep the probability of at least one new edit above 99%), we need to keep average expected new edits above five (see Fig.1D). To sustain editing throughout the typical *Drosophila* neuroblast lineage with 100 cell cycles, the Cas9 editing rate (*r*) and the number of Cas9 targets (*nU*) need to satisfy certain conditions (see Fig.1E). Using five as the minimal number of expected new edits per cell cycle, we determine the optimum editing rate to record 100 serial cell cycles to be 0.01 and the total required units to be 1353 (Fig.1E center). The efficiency of this scenario, derived as the ratio of the total units minimally required to cover the entire lineage (500) over the actual required number of units (1353), is 37%. Lowering the editing rate reduces the efficiency because the majority of units are left unedited by the end of 100 cell cycles (Fig.1E left). By contrast, a higher editing rate rapidly consumes excessive units and therefore would require astonishing numbers of total units (*e.g.* 1.7 million with rate of 0.1; Fig. 1E right) to sustain new edits throughout 100 serial cell cycles. The minimum number of required targets is attained when the editing rate is 0.01 for the case of 100 cell cycles, regardless of required confidence level (which determines the number of required targets) (Fig.1F). In general, the optimum editing rate is simply *r* = 1/*depth* (see Fig.1F legend).

Taken together, our modeling shows that the efficient use of targets highly depends on the editing rate.

### Cas9 base-editing occurs at near-optimum rate to encode 100 cell cycles

Given the importance of the editing rate in our modeling, it is critical to determine whether the actual rate of the Cas9 base-editor (BE2 modified by David Liu from BE1 to suppress base-excision repair[31]) makes these proposed experiments achievable. To test Cas9 base-editing rates *in-vivo*, specifically in the cycling neuroblasts of elongated *Drosophila* neuronal lineages, we engineered two UAS-GFP transgenes to detect C-to-T base-editing. We mutated the ATG start codon to ACG to silence GFP and also modified the surrounding sequences to create targets for two different gRNAs. Only in the presence of corresponding gRNAs can the Cas9 base-editor, BE2, elicit C-to-T mutation and thus restore the ATG start codon for GFP expression.

To monitor the C-to-T mutational dynamics in cycling neuroblasts, we drove the conditional UAS-GFP transgenes using dpn-GAL4 which selectively labels ~100 neuroblasts per brain lobe. Cas9 was driven by an actin promoter and gRNAs by U6 promoters. Despite ubiquitous induction of both Cas9 and the gRNA, we detected mosaic patterns of GFP activation in the developing CNS. This observation suggests limited Cas9 action prior to budding of neuroblasts from the neuroepithelium. We examined the GFP patterns at 2 days and 5 days after larval hatching (ALH). At 5d ALH, when larvae are easily sexed, it was clear that male (XY) brains contained more GFP positive neuroblasts than female (XX) brains. The Cas9 base-editor transgene resides on the X chromosome. Male flies compensate for having only one X chromosome by increasing the transcription level by approximately two-fold[32]. The increased number of GFP positive lineages in males is consistent with a stronger induction of the X-chromosome (containing the Cas9-BE2 transgene) in hemizygous males than heterozygous females. We therefore quantified male and female brains separately (Fig.2C and Supplementary Fig.1).

**Figure 2:**
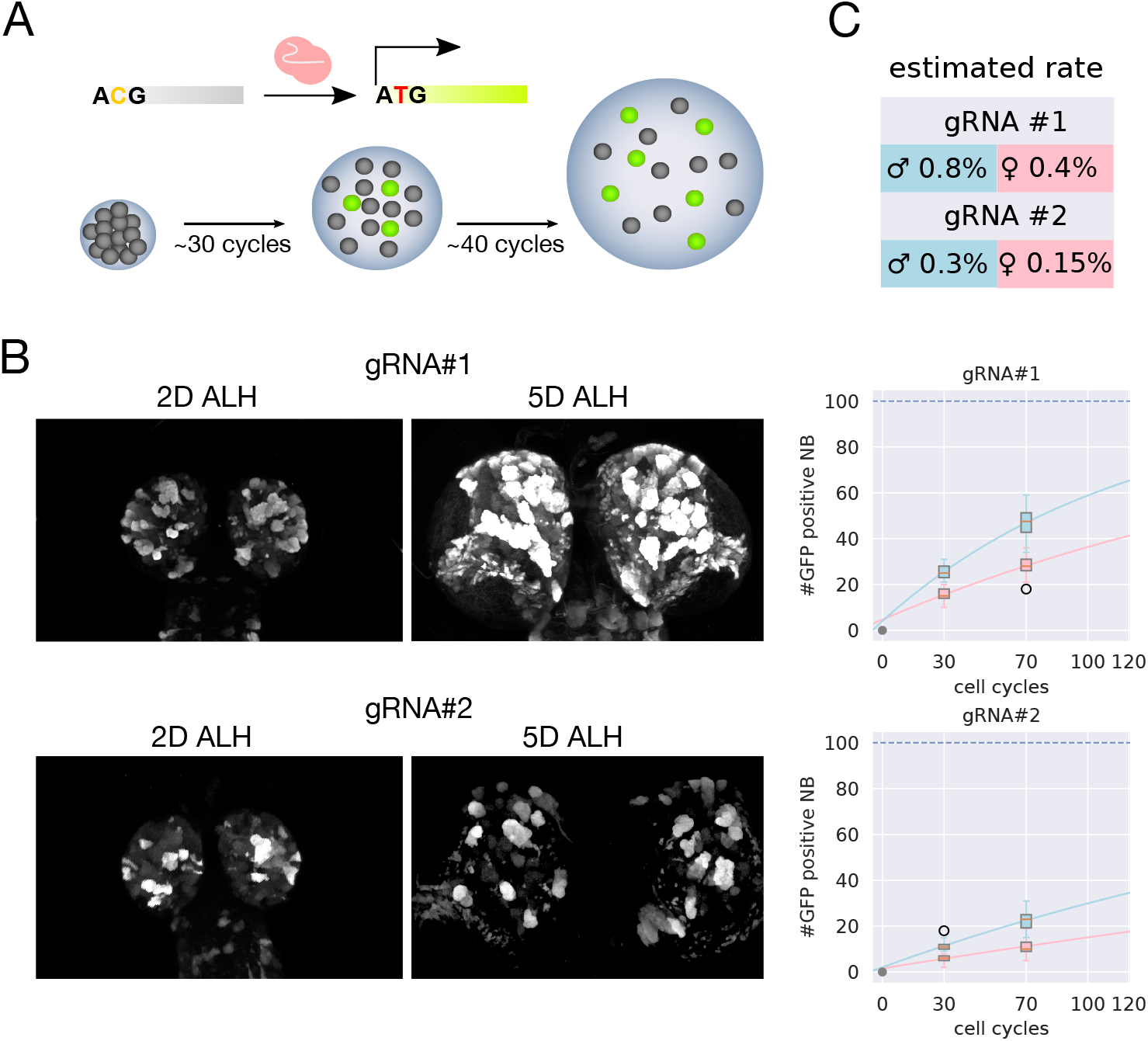
Estimation of Cas9 BE2 editing rate in *Drosophila* neuroblasts **A**: Experimental design to assay Cas9 BE2 editing rate *in-vivo* in *Drosophila* neuroblasts. GFP reporter with inactive start codon (ACG) produces no GFP flourescence (grey bar and grey circles). After C-to-T editing by Cas9 BE2, start codon becomes ATG and GFP is expressed (green bar and green circles). GFP positive neuroblasts (green circles) are counted at 2d ALH (~30 neuroblast divisions) and then at 5d ALH (~70 neuroblast divisions). **B**: Example confocal images for two gRNAs at 2d and 5d ALH. **C**: Fitted C-to-T editing rates for each gRNA. The sex of young larvae (2d ALH) was assigned via modeling with Gaussian mixture (Supplementary Fig.1). See Methods for details. n=136 for gRNA#1 and 156 for gRNA#2.

Following gross assessment, we counted GFP-positive neuroblasts in whole-mount larval brains imaged with confocal microscopy (Fig.2B). At 2d ALH the larval sex is not easily determined; therefore we assigned their sex subsequently by fitting a two-component Gaussian mixture model as a function of neuroblast count (Supplementary Fig.1). Plotting the number of fluorescent lineages revealed that neuroblasts undergoing C-to-T editing consistently increased from 2d to 5d ALH for all four gRNA/sex combinations (292 brain lobes in total). Using the cell cycle data of a fully mapped antennal lobe lineage[27], we assigned the cell cycle number at 2d ALH to be 30 and the cell cycle number at 5d ALH to be 70. We then used these numbers to calculate binomial success rates based on cell cycle numbers (Fig.2C). We found that the editing rates varied based on gRNA sequence—gRNA#1 is faster than gRNA#2. Also, as predicted by X chromosome dosage compensation, males have rates approximately double the rates of females. Encouragingly, the *in-vivo* editing rate of 0.84% detected in males with gRNA#1 is exceptionally close to the optimum rate to track 100 serial cell cycles (1%). This lends credence to our experimental plan. We therefore use 1% as the editing rate for all subsequent computer simulations of barcode generation.

### Combining multiple experiments compensates for limited coding units and cell/barcode loss

If the encoding is robust (*i.e.* there are always new edits at each cycle), then reconstruction of the lineage tree should be fairly easy. However, the minimum required targets to encode 100 cycles is 1353 for a confidence level of 99% (812 for 90%, see Fig.1F). This is a large number and not easily attainable *in-vivo*. Moreover, in practice, cell and barcode loss during experiments is unavoidable. Therefore, it is not possible with the currently available tools to reconstruct a full lineage tree from a single experiment. For these reasons, we next explore ways to combine multiple experiments.

To merge partial trees into a more complete tree, we need to know the identities of cells that are present in various subtrees. We can then establish correspondence of cells (leaf nodes) between partial trees and relate birth order information across trees built from different samples. In this section, we assume it is possible to distinguish individual cells based on gene expression profiles which are to be recovered simultaneously during barcode retrieval via single cell RNA-seq (see Discussion).

With known cell identities, we can in principle align partial trees to derive a complete tree. However, ordering errors are inevitable in single trees. For instance, due to Cas9 perdurance from neuroblasts, editing can occur in dividing GMCs and thus complicate the progeny ordering[33] based on the assumption that edits accumulate through serial stem-cell cell cycles. The objectives of merging partial trees should include improvements in accuracy and resolution in addition to complete coverage. With this intention, we conceived n-by-n matrices, termed order matrices, to describe pairwise order relationships across cells with distinguishable cell type information (Fig.3A-C). For a given set of clonally related cells, we determine the birth order among the recovered cells based on the numbers of edits present in the barcodes. We denote the value for a pairwise comparison as 1 if the cell in a row of the matrix carries fewer edits than the cell in a column and is thus deemed earlier born, or −1 if the converse. For undetermined or non-applicable situations (including tie, missing, or self-comparison), we assign a value of 0. The column-wise sum of the positive values results in a score for each cell type. If no errors exist (including inadequate encoding units and cell/barcode loss), the scores can be sorted to reveal the birth order. For clarity, we give very simple examples of scoring a single-tree order matrix (Fig.3B), and how the order matrices would appear with errors or missing data due to inadequate encoding units or cell/barcode loss (Fig.3C). In reality, a single tree order matrix will be far more complicated, containing multiple sources of error. In Fig.3D, we show a representative single-tree matrix with a *depth* of 50 and all categories of error combined. With each single-tree simulation, we can see the encoding process as well as the extent of random cell/barcode loss. Importantly, the editing rate optimized to minimize coding unit waste (calculated as 1/*depth*, Fig.1E) is also the optimum rate to retrieve order information (Fig.4A). This rate maximizes the chances of new edits occurring throughout the entire lineage.

**Figure 3:**
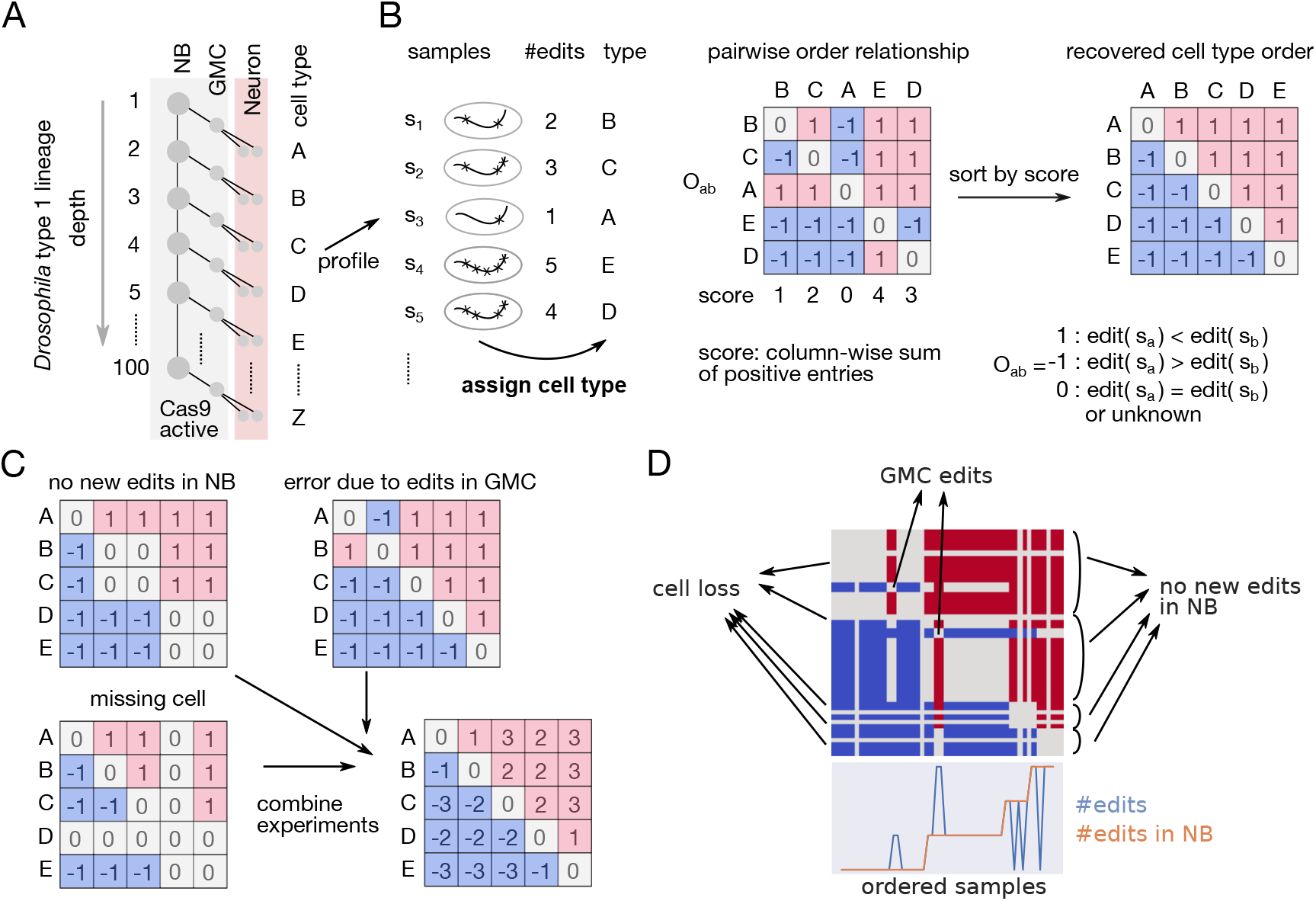
Recovering proper order from pairwise order relationships **A**: Schematics of *Drosophila* type 1 neuroblast lineage development. A special characteristic of the type 1 lineage is its linearity. Cas9 can be driven by *dpn* and assumed to be inactive in post-mitotic neurons. Neuronal cell-types are indicated with capital letters. Neurons are the target of simultaneous single cell RNA-seq and lineage barcode profiling. NB:neuroblast, GMC:ganglion mother cell. **B**: Schematics showing how to recover proper birth order when pairwise orders of recovered cell types are known. Left, samples *s_n_* are assigned with cell types and number of edits determined by RNA-seq. Center, cells are placed in a matrix and pairwise order relationship (given value of −1, 0, or 1) can be estimated from the number of edits. Right, cells are reordered according to the score (column wise sum of positive entries). **C**: Combining order matrices yields improved lineage order. Order matrices with various types of error are shown. Order information is absent (0) if there are no new edits in the neuroblast (NB) or if there is a missing cell or barcode sequence. Due to editing in GMCs, order estimated from number of edits can be swapped (blue in upper triangular region). To combine experiments, pairwise order information (−1, 0, or 1) are accumulated (simply summed) and the final order relationship is determined by comparing the counts of positive entries (red) across columns. **D**: An example of order relationships when there are mixtures of errors. GMC edits appear as blue in upper triangular region and red in lower triangle. Due to lack of edits, no ordering information can be obtained (gray region). Cell losses appear as vertical and horizontal zeros.

**Figure 4:**
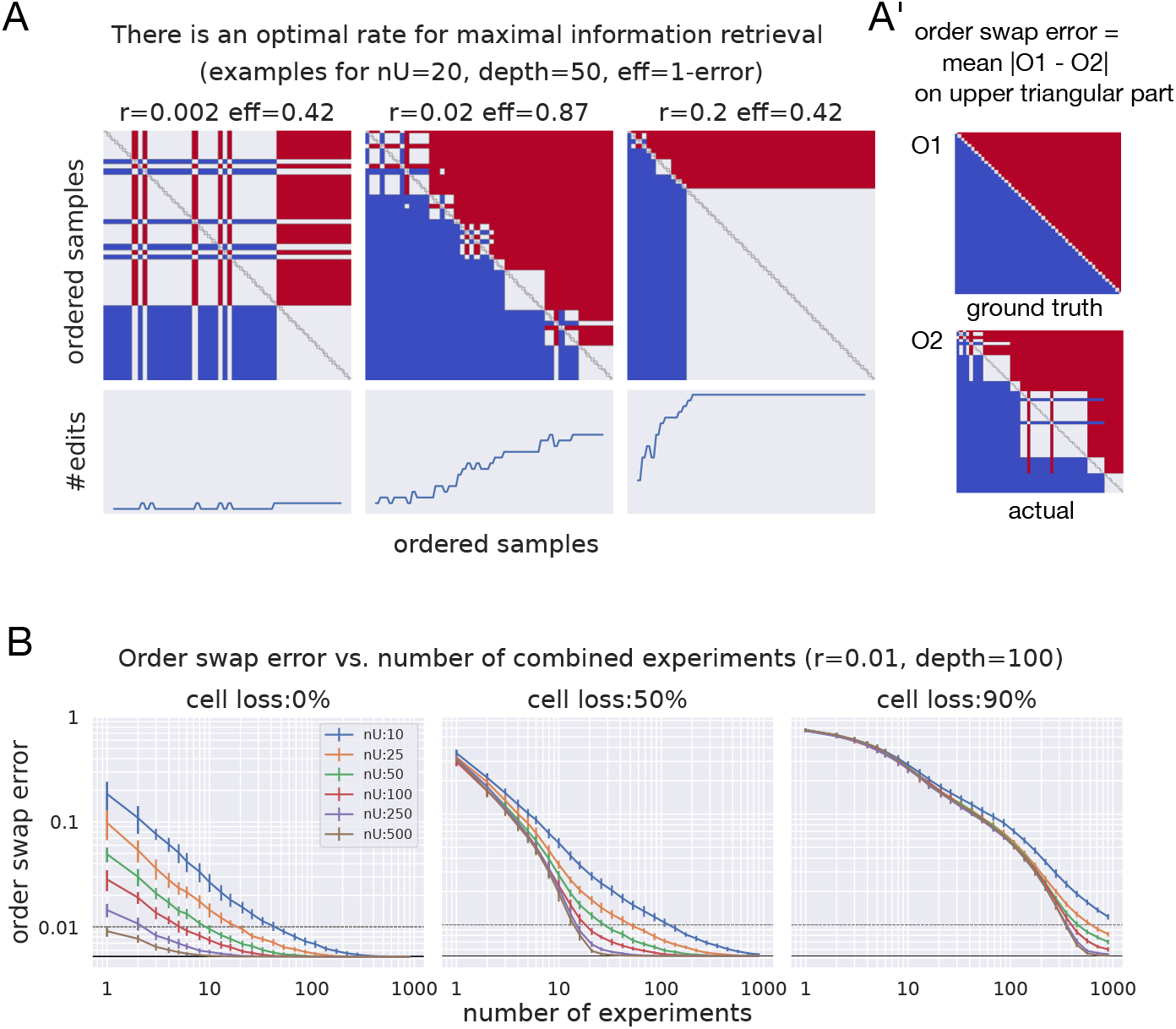
Combining experiments enables full lineage reconstruction despite low unit number and cell loss **A**: Examples of order information obtained by varying the editing rate. Samples (row and columns) are ordered according to developmental time. For simplicity, there is no cell loss. Top panels show order matrices; corresponding bottom panels show number of edits present in the ordered samples. Left, at a low rate (*r* = 0.002 = 0.1/*depth*), the unwanted GMC edits readily cause errors as the editing in NB occurs rarely (only near the end of the shown lineage). Center, *r* = 0.02 = 1/*depth* is the optimum rate to most efficiently obtain order information, as new edits are accumulated across the lineage. Right, at a rate much higher than optimum (*r* = 0.2 = 10/*depth*), all edits occur at the beginning of the lineage and no new information is retrieved afterwards. The optimum rate again can be easily derived from expected number of new edits at depth *d*: *n_d_* = *nUr*(1 − *r*)^*d*−1^. This time, *nU* is fixed and we want to maximize *n_d_*. Having more expected new edits at depth *d* ensures more expected new edits at the previous cycles and thus better order distinguishability. Efficiency (eff) is defined as 1 minus (order swap error). **A’**: Order swap error is the proportion of pairs with the wrong order relationship. **B**: Combining single-tree order matrices improves the accuracy of tree reconstruction. Barcodes with different numbers of coding units (*nU*) were simulated to encode a 100-depth linear lineage at the optimum editing rate of 1%. Simulations were further subjected to varying extent of random cell loss. Up to 1000 experiments were combined for each condition and this was repeated 20 times to derive the average error rate and standard deviation. Note the log scales for both number of cumulative experiments and order swap error in combined order matrices.

Such digital order matrices render the birth-order information additive across trees constructed from the same lineage. That is, individual single-tree order matrices with cells sorted in the same order can be summed up to derive combined scores (Fig.3C). We then score it by counting the red-shaded entries (*i.e.* positive values) for each column, and sort cells by these scores in the combined order matrix. To determine the accuracy of the combined order matrices, we quantified the deviation from the ground-truth order matrix by calculating “order swap error” as the mean of absolute value differences for those entries in the upper triangular part of the matrices (Fig.4A’).

To validate the above strategy for data pooling, we used simulations to determine the feasibility and number of experiments needed to achieve faithful reconstruction of 100 serial cell cycles under various conditions. We simulated lineages with barcodes consisting of 10 to 500 coding units (Fig.4B left). For each condition, we additionally simulated cell/barcode loss of 50% (Fig.4B, center) and 90% (Fig.4B, right). Using order swap error as a readout of accuracy, we monitored how combining order matrices (increasing numbers of experiments) cumulatively improves lineage reconstruction (Fig.4B). In the conditions with 50% cell loss, we find it feasible to reconstruct the entire 100-depth linear lineage with an order swap error of below 1% by combining just tens of experiments. Further, it is remarkable that even barcodes with only 10 coding units permit faithful reconstruction of 100 serial cell cycles, although several hundred experiments need to be combined. As to severe cell/barcode loss (90%), it takes many more experiments to restore an entire lineage. Also, the number of coding units becomes relevant only after combining 200+ experiments, indicating that the effect of cell loss dominates the error rate when the number of experiments is relatively small.

In summary, with order matrices, we can effectively combine multiple experiments to compensate for inadequate number of coding units or cell/barcode loss and thus realize full lineage reconstruction.

### Effects of cellular redundancy and cell-type recurrence on lineage reconstruction

In the previous section, we assume that post-mitotic cells are individually unique and can be matched across experiments (resulting in 200 individual cells). However, cellular redundancy can exist with multiple consecutively derived cells belonging to a single cell type. The known degrees of redundancy vary from yielding only seven neuron types through ~300 serial cell cycles[34] to producing over 40 morphologically distinct types via ~100 asymmetric divisions[27, 28]. In such cases, the order matrices should deal with cell types rather than individual cells. This reduces temporal complexity and likely also moderates challenges in lineage reconstruction. To model cell type redundancy, we simulated lineages that undergo 100 serial cell cycles but yield fewer cell types. As expected, reducing the temporal complexity greatly eased the reconstruction of protracted linear lineages (Fig.5A). Despite 90% cell loss, one can faithfully map 50 serial cell types (with 4 cells per type) by combining just ~40 experiments. Notably, cellular redundancy also diminishes the need for copious coding units, as increasing coding units from 50 to 500 provides only marginal benefits.

**Figure 5:**
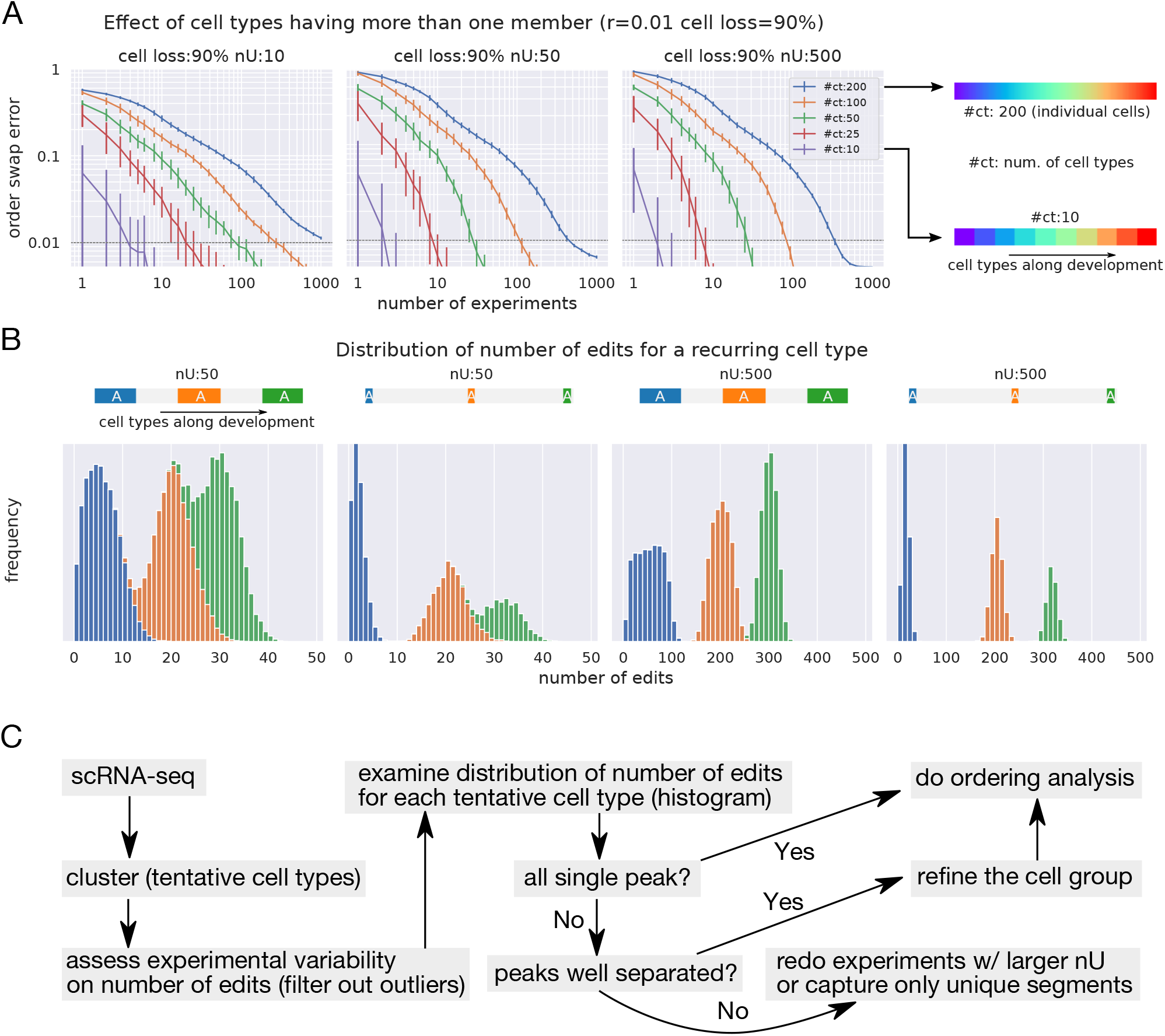
Effect of cell redundancy and recurrence **A**: Cellular redundancy, or the same cell type born in a given temporal window, allows for the grouping of cells in a consecutive manner along developmental time. Grouping in this manner effectively reduces the temporal complexity and decreases the required number of experiments to recover full sequence order, as revealed by the plots of order swap error vs. number of combined experiments (mean of 20 simulations +/− standard deviation). Depth of lineage:100, edit rate:1%. All experiments have 90% cell loss and the different line color indicate varing number of coding units (*nU*). **B**: When one cell group (*e.g.* cell type A) arises from multiple windows (colored bars), plotting the distribution of the number of edits from multiple experiments (here 1000) can determine whether the recurrent cells can be separated into discrete temporal subsets for birth order analysis. Histograms show distributions of edit numbers under different conditions, varying both the number of coding units (*nU*) and the duration of the windows yielding the same cell type. For clarity, the histograms of edit numbers are differently colored for different windows, but would not be distinguishable in an experimental setting. **C**: Proposed workflow for dynamic lineage encoding to achieve synergistic cell type and cell lineage mapping.

However, it is also possible that indistinguishable cells may arise from multiple windows in a given lineage. This could be due either to recurrent production of the same cell types[28] or technical challenges in distinguishing closely related cell types. In either case, we need to avoid clustering temporally distinct cells into a single group for birth order analysis. One way to achieve this is to separate groups based on the numbers of cumulative edits, as edits should increase along lineage depth. Plotting the distribution of edits of a single distinguishable cell type from several experiments can reveal the presence of recurrent cell types (Fig.5B). Both the duration as well as the position of the cell-production windows govern the extent of overlap in the distribution of edit numbers, and thus how well they can be separated for order analysis. Further, as observed from simulations comparing 50 and 500-unit barcodes, increasing the number of coding units can greatly narrow the distribution of edits, facilitating identification of recurrent cell types prior to lineage reconstruction (Fig.5B).

In conclusion, we have established a strategy to map serially derived cell types in linear elongated lineages with dynamic barcodes. We provide the following workflow (Fig.5C) to ensure accuracy of data before performing order analysis: First, we perform single cell RNA-seq and barcode sequencing of multiple lineage tracing experiments for a given lineage. We then cluster distinguishable cell types and determine if there are any experiments that are outliers (issues with normal progression of editing). We then assess the lineage for signs of recurrent cell types by plotting the distribution of edits of each distinguishable cell type. Only cell types with single-peak distribution of edit numbers can be directly subject to birth order analysis using order matrices. For those with multiple peaks but narrow distribution, the cell types need to be classified discretely before order analysis. For cell types exhibiting broad/overlapping distribution, it is important to resolve them into distinct temporal groups prior to order analysis (see Discussion). In this manner, we can map protracted *Drosophila* type I neuronal lineages with a modest number of barcodes (*nU* = 50) despite substantial cell/barcode loss and the presence of recurrent cell types.

## Discussion

In this paper, we have examined the requirements to apply CRISPR-based lineage tracing combined with single cell RNA-seq on an prolonged stem-cell lineage. We chose to model conditions using a Cas9 base-editor (see below). We found that the optimum editing rate (*r*) for efficient use of targets depends on the number of cell cycles to be tracked *r* = 1/*depth*. In the case of a typical type I neuronal lineage in the *Drosophila* central brain, which undergoes 100 cell cycles, the optimum Cas9 editing rate is 1% (*i.e.* editing 1% of available Cas9 targets per cell cycle on average). We measured the *in-vivo* editing rate of a Cas9 base-editor (BE2 modified by David Liu from BE1 to suppress base-excision repair[31]) to edit a single target in *Drosophila* neural stem cells. We found editing rates (0.15-0.84%) can be reasonably close to the optimum rate of 1%. This suggests that performing CRISPR-based lineage tracing with the Cas9 base-editor is feasible for these extremely deep lineages. Still, for the case of an ideal single experiment, the required number of CRISPR targets (*nU*) is quite large (1353 to have a 99% chance of editing at the 100th cell cycle). To circumvent this issue, we utilized the simple linear tree structure of type I lineages to easily combine information from multiple experiments. This substantially reduces the required number of targets, thus permitting dense reconstruction despite low target numbers. Nevertheless, the number of required experiments needs to be high enough to accommodate for the loss of cells/barcode sequences which can be substantial with single-cell RNA-seq. Additionally, the number of cell types produced changes the number of experiments needed. We estimate that with 50 coding units and 90% cell/barcode loss, we would need approximately 50 experiments (individual flies) to resolve a lineage that generates 50 distinct cell types during its 100 divisions. This is experimentally achievable, assuming both the ability to target/identify lineage(s) of interest and ideally the ability to identify individual organisms for multiplexed lineage tracing.

### Advantages of Cas9 base-editors

Cas9 base-editors have multiple potential advantages over Cas9 nuclease in the encoding of serial cell cycles with cumulative independent mutations. Avoiding double strand break would permit a high number of arrayed targets without concerns over inter-target deletion. Inter-target deletions caused by Cas9 nuclease[10], can drastically reduce the coding capacity and simultaneously erase earlier edits[30]. These aspects of Cas9 nuclease are particularly detrimental to the tracking of deep linear lineages. By contrast, base-editors independently edit individual targets. One can thus pack many targets into a single transgene, to increase target numbers while facilitating barcode retrieval. As for edit outcomes, indels from Cas9 nuclease are less predictable and more variable than base editing. However, Cas9 cytidine deaminase can elicit C-to-T mutation at multiple sites within a 5-bp window of the target. By placing two Cs within the 5-bp window, one can generate three possible outcomes per target[35]. Further, some Cas9 base-editors are able to edit all four bases (ATGC) and thus create more outcomes[36]. Nonetheless, to be conservative, we set the number of edit outcomes as two (*nL* = 2) in our modeling. This does not appear to limit our ability to track 100 serial cell cycles. In fact, we have previously observed that the number of alleles per unit is much less important compared to the rate of editing and the number of coding units[37, 38]. A recent study on cell phylogenic analysis has also concluded similarly about the Cas9 base-editor’s advantage in having many more targets despite limited edit outcomes[39].

While the number of targets affect coding density, the editing rate determines the pattern of unit usage over serial cell cycles. The optimum rate of 1/*depth* ensures encoding with the highest efficiency. To track *Drosophila* type I neuronal lineages with 100 cell cycles, a Cas9 base-editor with a relatively low editing rate is favored over Cas9 nuclease. An estimated rate for Cas9 nuclease in *Drosophila* embryogenesis is ~12% per cell cycle[30], much higher than the desired rate of 1%. By contrast, the editing rate of the likely more potent BE3 (compared to BE2) in cultured cells could be ~2%[29]. Despite some shortcomings in our rate evaluation in fly neural stem cells, we found the rates of BE2 to be lower than 1% by relatively small margins. As the editing rate was dependent upon the target/gRNA sequence (see Fig.2), this rate can likely be optimized. For lineage tracing experiments, we would drive the Cas9 base-editor via a binary induction system (rather than directly under an actin promoter) to restrict edits to cycling progenitors. The more versatile control of Cas9 base-editor expression by binary induction should facilitate optimization of editing rates. Binary induction should also eliminate the observed gender-dependent 2-fold difference in the editing rate, which is likely due to dosage compensation of the X-chromosome containing the BE2 transgene. In sum, the Cas9 base-editor stands out as a better choice to track protracted neuronal lineages with dynamic barcodes.

### Combining experiments to provide identifiable cell types with birth-order information

Combining data from multiple CRISPR-based dynamic lineage tracing experiments is essential for a comprehensive understanding of a given lineage. From a single experiment, lineage reconstruction can yield only a partial tree. This is due to sparse cell sampling which results in an insufficient number of unique leaves for full tree construction and/or suboptimum encoding process. To derive a complete tree, therefore requires multiple sampled lineages. Unfortunately, samples from different experiments cannot be directly combined, as lineage barcode generation is unique to each animal.

In phylogeny field, combining multiple partial trees have been accomplished using either supermatrix approach (concatenation of individual evidences, in this case barcodes from different experiments) or supertree approach (finding a tree that is most consistent with triplets or quartes partial trees from individual trees)[40, 41, 42]. However, these approaches have not been applied to the CRISPR-lineage tracing field. In this article, rather than applying the methods from traditional phylogeny field, we took advantage of the linear topology of *Drosophila* type 1 lineages and devised a way to use pairwise orderings of the cells to combine multiple experiments. We use the number of CRISPR edits to estimate the pairwise order in individual experiment. To combine information from multiple experiments, we accumulated pairwise order information from each experiment into a more complete order information. This process improves both accuracy and completeness. By combining experiments, we found that it is feasible with a fairly low number of targets (*nU* = 50) to resolve a 100 cell-cycle lineage with 50 unique cell types (Fig. 5A), provided there are no recurring cell types (Fig. 5B). Here, we dealt with simple linear lineages to combine experiments. However, the same idea can be readily extended to more general cases by using triplet ordering of cells instead of pairwise ordering (Sugino and Lee in preparation).

To combine order relationships across experiments, the ability to match corresponding cell types across experiments is essential. Despite unavoidable dropouts with current single cell RNA-seq methods, massive analysis of single-cell transcriptomics permits identification of cell types with unique marker combinations. One can then assign the profiled single cells with identities based on the combination of markers, even allowing for some dropouts. This strategy works effectively even among closely related neuron types made by a common progenitor. For instance, an analysis of ~600 single-cell transcriptomes profiled from a Drosophila antennal lobe neuronal lineage comprised of 27 morphological types revealed 18 clusters with distinguishable gene expression patterns[43].

### Managing cell-type redundancy and recurrence

The number of experiments needed for accurate lineage mapping depends upon the number of unique cell types within a lineage (See Fig. 5). In cases where the transcriptomic cell types are not yet determined, we expect to identify tentative cell types through the initial analysis of pooled single cell RNA-seq data. We then need to evaluate if each identifiable cell type arises from the same approximate developmental window in all samples. This can be addressed by examining the distribution patterns of edit numbers (see Fig. 5B). Only cell types with either single peaks or multiple peaks that are clearly defined can be included for collective order matrix analysis. Redundant cell types (i.e. a block of a particular cell type) would present as a broad peak but would be clearly separable from other distributions.

To include cell types in a collective order matrix that do not have well-defined peaks, overlapping peaks must first be separated. To separate peaks for cell types with broad, potentially mixed distributions, it may be reasonable to resolve the cell production window(s) by increasing the number of coding units. Alternatively, one can target discrete temporal windows for stage-specific lineage tracing via temporal induction of Cas9[15, 44]. This partial lineage tracing should separate otherwise indistinguishable cells based on the timing of their birth. Cells born in different windows but assigned to a single transcriptomic type can be scrutinized further to uncover possible subtypes with subtle differences in gene expression

### Unique identifier labeling for multiplexed lineage tracing

With the availability of a method to combine multiple experiments, the scalability (i.e. how many lineages can be simultaneously traced and profiled) becomes a major limiting factor. By combining approximately 50 experiments, it should be feasible to resolve 50 cell types in a 100 cell-cycle neuronal lineage, provided cell types do not recur (Fig. 5). Although 50 flies (needed in the above case) can be individually sorted and libraries can be constructed separately, this would be both time consuming and expensive. For scalability, pooling animals would be ideal. This could be achieved if individual animals (if tracking selected lineages) or even individual neuroblast lineages (if tracking many/all lineages) could be distinguished genetically.

Orthogonal CRISPR systems can be utilized for independent organismal or neuroblast marking in conjunction with dynamic lineage encoding. Orthogonal CRISPR systems include different forms of Cas9[45] or Cas9 nucleases that recognize different target sequences based on their PAMs. To create animal or neuroblast unique identifiers (UID), Cas9 nuclease is superior to Cas9 base-editor, as Cas9 nuclease achieves higher efficiency and more diverse outcomes per site. To give each animal a UID, Cas9 nuclease can be expressed in post-mitotic germ cells of parental flies to create diverse offspring-specific genetic scars. To bestow neuroblast UIDs, the Cas9 variant should be acutely and robustly induced just before the start of neuroblast divisions to create distinct lineage-specific genetic scars. If we use a separate array of 50 targets and on average 10% of the targets are edited (5 out of 50), then in theory, we would have over 2-billion different possible UIDs (assuming *nL* = 4), which is more than adequate to distinguish the 2 (hemispheres) × 100 (type I neuroblasts) × 50 (flies)=10,000 neuroblast lineages.

### Conclusion

Dynamic lineage tracing facilitates the simultaneous and synergistic mapping of transcriptomic cell types and cell lineages. Here, we demonstrate a strategy to combine pairwise order information across samples to merge partial lineage trees into a complete tree. We determined relevant experimental parameters and found it possible to achieve dense reconstruction of *Drosophila* type I neuronal lineages with modest number of barcodes. We have also extended this approach to more general (non-linear) trees using triplet orderings of cells instead of paired orderings (Sugino & Lee, manuscript in preparation).

## Materials and Methods

### Simulation of barcode generation

Stochastic behavior of the CRISPR barcode is generated using NumPy (http://numpy.org) random number generator (numpy.random.uniform). The Python source code and associated Jupyter notebooks are deposited in GitHub (http://github.com/kensugino/drosophila_type1_lineage_barcode).

### Analysis software

The following programs and libraries are used for the analysis and figure generation: Python (http://www.python.org), matplotlib[46], pandas[47], sklearn[48], seaborn (http://seaborn.pydata.org), jupyter (http://jupyter.org).

### Cas9 reporter experiments

The gRNAs used in Fig.2 were chosen based on the following criteria: their target should comprise the sequence ACG within the base-editor mutational window (3-10bp of the protospacer). High predicted on-target and off-target scores using the CRISPR tool on Benchling (https://benchling.com). For base-editor Cas9 targets, on-target scores >10 and off-target scores >90 were predicted for gRNAs that worked efficiently in our hands. Scores: gRNA#1 (on-target: 22; off-target: 99.1) and gRNA#2 (on-target: 19; off-target: 98.1).

UAS-GFP base-editing reporters were constructed by replacing the ATG start codon with the target for gRNA#1 or gRNA#2 and rendering the to-be-edited ACG in the same reading frame as GFP. The corresponding U6-gRNA was placed in the same construct with the UAS-reporter for site-specific germline transformation. Male flies carrying the reporter/gRNA transgene, together with dpn-GAL4 were crossed with female flies carrying actin-BE2 on the X chromosome. Synchronized larvae carrying all above transgenes were dissected for analysis of GFP patterns in developing brains.

### Estimation of editing rate from counts data

The editing rate is estimated from the count data as follows. The units remaining at 30 cycle (*u*_30_) is the original number of units (total number of neuroblast *u*_0_ = 100) minus edited neuroblast at 30 cycle (*e*_30_), *u*_30_ = *u*_0_ − *e*_30_. According to the binomial model, the number of additional edits after 40 cell cycle is *e*_70_ − *e*_30_ = *u*_30_(1 − *s*^40^), where *s* = 1 − *r*. From this, we have *s* = (1 − (*e*_70_ − *e*_30_)/*u*_30_)^1/40^ and the editing rate *r* = 1 − *s*. Thus, the rate is calculated solely from the behavior in between 30th cycle and 70th cycle.

To obtain the origin (the presumed start of editing assuming the editing rate were the same as in between 30th and 70th cycle), we denote the origin *x*_0_ and used 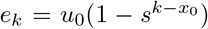, where *k* = 30, 70. From this, 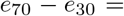 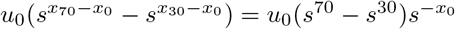 and therefore, *x*_0_ = *log*(*u*_0_(*s*^70^ − *s*^30^)/(*e*_70_ − *e*_30_))/*log*(*s*).

## Acknowledgments

We thank Jorge Garcia-Marques for critical reading and comments.

## Supplements

## SuppFig1 for Figure 2: separating 2D ALH samples into two groups

**Figure Supp.1:**
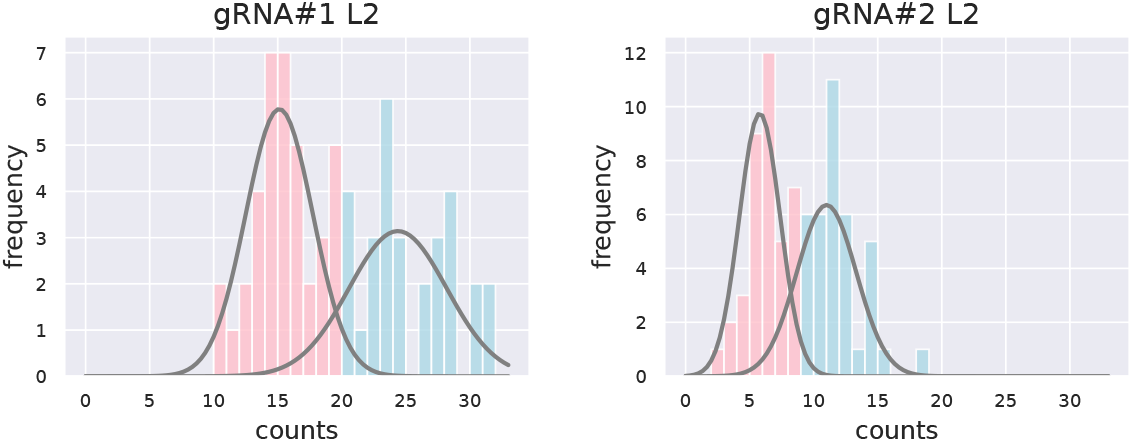
Separating male and female brains at 2d ALH. We observed clear separation of GFP+ neuroblast counts between male and female brains at 5d ALH (Fig.2C). This suggests sex dependent differences in Cas9 editing rate (likely due to differential expression of the Cas9-BE2 transgene location on the X chromosome). For proper estimation of the editing rate, it is important to calculate the rate separately in males and females. However, at 2d ALH, male and female larvae are not easily distinguishable. Plotting the counts of GFP+ neuroblasts at 2d ALH in a histogram (for each gRNA) shows two separate populations. We, therefore, fit the counts at 2d ALH with a Gaussian mixture model and designated the population with higher counts as males and the population with lower counts as females. These separation results were used to estimate the editing rate (Fig.2C).

## Notes

### Competing Interest Statement

The authors have declared no competing interest.

## References

[1] Ken Sugino et al. “Mapping the Transcriptional Diversity of Genetically and Anatomically Defined Cell Populations in the Mouse Brain”. In: eLife 8 (Apr. 12, 2019). Ed. by Catherine Dulac, Chris P Ponting, and Nenad Sestan, e38619.

[2] Raymond E. Keller. “Vital Dye Mapping of the Gastrula and Neurula of Xenopus Laevis: I. Prospective Areas and Morphogenetic Movements of the Superficial Layer”. In: Developmental Biology 42.2 (Feb. 1, 1975), pp. 222–241.

[3] U. Deppe et al. “Cell Lineages of the Embryo of the Nematode Caenorhabditis Elegans”. In: Proceedings of the National Academy of Sciences 75.1 (Jan. 1, 1978), pp. 376–380.

[4] J. E. Sulston et al. “The Embryonic Cell Lineage of the Nematode Caenorhabditis Elegans”. In: Developmental Biology 100.1 (Nov. 1983), pp. 64–119.

[5] Charles B. Kimmel and Robert D. Law. “Cell Lineage of Zebrafish Blastomeres: III. Clonal Analyses of the Blastula and Gastrula Stages”. In: Developmental Biology 108.1 (Mar. 1, 1985), pp. 94–101.

[6] C. Walsh and C. L. Cepko. “Widespread Dispersion of Neuronal Clones across Functional Regions of the Cerebral Cortex”. In: Science 255.5043 (Jan. 24, 1992), pp. 434–440.

[7] Dawn L. Zinyk et al. “Fate Mapping of the Mouse Midbrain–Hindbrain Constriction Using a Site-Specific Recombination System”. In: Current Biology 8.11 (May 21, 1998), pp. 665–672.

[8] Liqun Luo. “Fly MARCM and Mouse MADM: Genetic Methods of Labeling and Manipulating Single Neurons”. In: Brain Research Reviews 55.2 (Oct. 2007), pp. 220–227.

[9] Rong Lu et al. “Tracking Single Hematopoietic Stem Cells in Vivo Using High-Throughput Sequencing in Conjunction with Viral Genetic Barcoding”. In: Nature Biotechnology 29.10 (10 Oct. 2011), pp. 928–933.

[10] Aaron McKenna et al. “Whole-Organism Lineage Tracing by Combinatorial and Cumulative Genome Editing”. In: Science 353.6298 (July 29, 2016), aaf7907.

[11] Samuel D. Perli, Cheryl H. Cui, and Timothy K. Lu. “Continuous Genetic Recording with Self-Targeting CRISPR-Cas in Human Cells”. In: Science 353.6304 (Sept. 9, 2016).

[12] Stephanie Tzouanas Schmidt et al. “Quantitative Analysis of Synthetic Cell Lineage Tracing Using Nuclease Barcoding”. In: ACS Synthetic Biology 6.6 (June 16, 2017), pp. 936–942.

[13] Reza Kalhor et al. “Developmental Barcoding of Whole Mouse via Homing CRISPR”. In: Science 361.6405 (Aug. 31, 2018).

[14] Anna Alemany et al. “Whole-Organism Clone Tracing Using Single-Cell Sequencing”. In: Nature 556.7699 (Apr. 2018), pp. 108–112.

[15] Bushra Raj et al. “Simultaneous Single-Cell Profiling of Lineages and Cell Types in the Vertebrate Brain”. In: Nature Biotechnology 36.5 (May 2018), pp. 442–450.

[16] Bastiaan Spanjaard et al. “Simultaneous Lineage Tracing and Cell-Type Identification Using CRISPR-Cas9-Induced Genetic Scars”. In: Nature Biotechnology 36.5 (May 2018), pp. 469–473.

[17] Michelle M. Chan et al. “Molecular Recording of Mammalian Embryogenesis”. In: Nature 570.7759 (7759 June 2019), pp. 77–82.

[18] Isabel Espinosa-Medina et al. “High-Throughput Dense Reconstruction of Cell Lineages”. In: Open Biology 9.12 (Dec. 2019), p. 190229.

[19] Aaron McKenna and James A. Gagnon. “Recording Development with Single Cell Dynamic Lineage Tracing”. In: Development 146.12 (June 15, 2019).

[20] Nanami Masuyama, Hideto Mori, and Nozomu Yachie. “DNA Barcodes Evolve for High-Resolution Cell Lineage Tracing”. In: Current Opinion in Chemical Biology. Synthetic Biology • Synthetic Biomolecules 52 (Oct. 1, 2019), pp. 63–71.

[21] H. Lin and T. Schagat. “Neuroblasts: A Model for the Asymmetric Division of Stem Cells”. In: Trends in genetics: TIG 13.1 (Jan. 1997), pp. 33–39.

[22] Catarina C. F. Homem and Juergen A. Knoblich. “Drosophila Neuroblasts: A Model for Stem Cell Biology”. In: Development (Cambridge, England) 139.23 (Dec. 1, 2012), pp. 4297–4310.

[23] Gerhard M. Technau, Christian Berger, and Rolf Urbach. “Generation of Cell Diversity and Segmental Pattern in the Embryonic Central Nervous System of Drosophila”. In: Developmental Dynamics 235.4 (2006), pp. 861–869.

[24] Chris Q. Doe. “Temporal Patterning in the Drosophila CNS”. In: Annual Review of Cell and Developmental Biology 33 (Oct. 6, 2017), pp. 219–240.

[25] Rosa Linda Miyares and Tzumin Lee. “Temporal Control of Drosophila Central Nervous System Development”. In: Current Opinion in Neurobiology 56 (June 2019), pp. 24–32.

[26] T. Lee, A. Lee, and L. Luo. “Development of the Drosophila Mushroom Bodies: Sequential Generation of Three Distinct Types of Neurons from a Neuroblast”. In: Development (Cambridge, England) 126.18 (Sept. 1999), pp. 4065–4076.

[27] Hung-Hsiang Yu et al. “A Complete Developmental Sequence of a Drosophila Neuronal Lineage as Revealed by Twin-Spot MARCM”. In: PLoS biology 8.8 (Aug. 24, 2010).

[28] Ying-Jou Lee et al. “Conservation and Divergence of Related Neuronal Lineages in the Drosophila Central Brain”. In: eLife 9 (Apr. 7, 2020).

[29] Byungjin Hwang et al. “Lineage Tracing Using a Cas9-Deaminase Barcoding System Targeting Endogenous L1 Elements”. In: Nature Communications 10.1 (1 Mar. 15, 2019), p. 1234.

[30] Irepan Salvador-Martínez et al. “Is It Possible to Reconstruct an Accurate Cell Lineage Using CRISPR Recorders?” In: eLife 8 (Jan. 28, 2019). Ed. by Aviv Regev and Patrick Hsu, e40292.

[31] Alexis C. Komor et al. “Programmable Editing of a Target Base in Genomic DNA without Double-Stranded DNA Cleavage”. In: Nature 533.7603 (May 2016), pp. 420–424.

[32] Marnie E. Gelbart and Mitzi I. Kuroda. “Drosophila Dosage Compensation: A Complex Voyage to the X Chromosome”. In: Development 136.9 (May 1, 2009), pp. 1399–1410.

[33] Jorge Garcia-Marques et al. “Unlimited Genetic Switches for Cell-Type-Specific Manipulation”. In: Neuron 104.2 (Oct. 23, 2019), 227–238.e7.

[34] Yoshinori Aso et al. “The Neuronal Architecture of the Mushroom Body Provides a Logic for Associative Learning”. In: eLife 3 (Dec. 23, 2014), e04577.

[35] Holly A. Rees and David R. Liu. “Base Editing: Precision Chemistry on the Genome and Transcriptome of Living Cells”. In: Nature reviews. Genetics 19.12 (Dec. 2018), pp. 770–788.

[36] Rina C. Sakata et al. “A Single CRISPR Base Editor to Induce Simultaneous C-to-T and A-to-G Mutations”. In: bioRxiv (Aug. 8, 2019), p. 729269.

[37] Ken Sugino et al. “Theoretical Modeling on CRISPR-Coded Cell Lineages: Efficient Encoding and Optimal Reconstruction”. In: bioRxiv (Apr. 14, 2019), p. 538488.

[38] Ken Sugino and Tzumin Lee. “Robust Reconstruction of CRISPR and Tumor Lineage Using Depth Metrics”. In: bioRxiv (Apr. 15, 2019), p. 609107.

[39] Matthew G. Jones et al. “Inference of Single-Cell Phylogenies from Lineage Tracing Data Using Cassiopeia”. In: Genome Biology 21.1 (Apr. 14, 2020), p. 92.

[40] Bernard R. Baum. “Combining Trees as a Way of Combining Data Sets for Phylogenetic Inference, and the Desirability of Combining Gene Trees”. In: TAXON 41.1 (1992), pp. 3–10.

[41] M. A. Ragan. “Phylogenetic Inference Based on Matrix Representation of Trees”. In: Molecular Phylogenetics and Evolution 1.1 (Mar. 1992), pp. 53–58.

[42] Michael J. Sanderson, Andy Purvis, and Chris Henze. “Phylogenetic Supertrees: Assembling the Trees of Life”. In: Trends in Ecology & Evolution 13.3 (Mar. 1, 1998), pp. 105–109.

[43] Hongjie Li et al. “Classifying Drosophila Olfactory Projection Neuron Subtypes by Single-Cell RNA Sequencing”. In: Cell 171.5 (Nov. 16, 2017), 1206–1220.e22.

[44] Sarah Bowling et al. “An Engineered CRISPR-Cas9 Mouse Line for Simultaneous Readout of Lineage Histories and Gene Expression Profiles in Single Cells”. In: Cell 181.6 (June 11, 2020), 1410–1422.e27.

[45] Kevin M. Esvelt et al. “Orthogonal Cas9 Proteins for RNA-Guided Gene Regulation and Editing”. In: Nature Methods 10.11 (Nov. 2013), pp. 1116–1121.

[46] J. D. Hunter. “Matplotlib: A 2D Graphics Environment”. In: Computing in Science & Engineering 9.3 (2007), pp. 90–95.

[47] Wes McKinney. “Data Structures for Statistical Computing in Python”. In: Proceedings of the 9th Python in Science Conference. Ed. by Stéfan van der Walt and Jarrod Millman. 2010, pp. 51–56.

[48] F. Pedregosa et al. “Scikit-Learn: Machine Learning in Python”. In: Journal of Machine Learning Research 12 (2011), pp. 2825–2830.

